# Intraspecific differences in alkaline tolerance in brook stickleback (*Culaea inconstans*) inhabiting neutral and alkaline lakes

**DOI:** 10.1101/2024.11.30.626187

**Authors:** Alex M. Zimmer, Charles Bernard, Marina Giacomin, Anne-Marie Dion-Côté, Greg G. Goss, Chris N. Glover

**Author notes:** Corresponding author: Alex M. Zimmer, Department of Biological Sciences, 355 Campus Ring Road, PO Box 5050, Saint John, NB, Canada E2L 4L5.

## Abstract

Exposure to alkaline water (pH > 9.0) is physiologically challenging for fish, yet our understanding of the physiology of alkaline tolerance in fishes is limited to a small number of ihighly specialized species. This study aimed to characterize mechanisms of alkaline tolerance in brook stickleback (*Culaea inconstans*), a fish species with a broad pH habitat range, including highly alkaline waters such as Buffalo Lake (pH = 9.2) in Alberta, Canada. Stickleback from Buffalo Lake and a neutral reference lake (Buck Lake; pH = 8.2) were collected from the wild and acclimated to common conditions (pH = 8.0) for at least 2 months. Both populations were then exposed to alkaline conditions (pH = 9.5), resulting in a significant decrease in survival (14% by 7 d of exposure) in Buck Lake fish, but no mortality in Buffalo Lake stickleback. In a 4-d exposure to alkaline water, fish from both populations experienced characteristic inhibitions of ammonia excretion followed by subsequent recovery, in conjunction with an accumulation of ammonia within the body. However, no differences were observed between populations. Analysis of tissue Na^+^ and Cl^-^ content showed a more pronounced decrease in Cl^-^ in Buck Lake fish, suggesting that tighter regulation of Cl^-^ homeostasis and/or acid-base balance may be an important feature of alkaline tolerance. RNA-sequencing analysis highlighted large differences in gene expression between the alkaline and neutral lake populations, and in response to alkaline exposure. Few of these changes in the expression involved genes known to be associated with nitrogen, ion, or acid-base balance. These data indicate that alkaline tolerance is higher in brook stickleback resident to an alkaline lake than those sourced from a neutral lake, a trait that may be related to differences in physiological and transcriptomic responses to alkaline exposure.

## Introduction

(Brauner et al., 2013; Danulat, 1995; Wilkie and Wood, 1996; Wood, 2022)(Pinheiro et al., 2021)(Danulat, 1995; Wood, 2022)Alkaline waters are generally considered extremely challenging to fish, with exposure to elevated pH (pH > 9.0) resulting in disturbances in nitrogen, acid-base, and ion balance. In most fishes, 80-90% of nitrogenous waste arising from protein catabolism is excreted across the gills as ammonia, with urea accounting for the majority of the remaining proportion (Wood, 1993). The current consensus is that ammonia excretion occurs as facilitated NH_3_ diffusion and that this process relies on an outwardly directed NH_3_ partial pressure (P_NH3_) gradient that is maintained by an acid-trapping mechanism in the apical gill boundary layer whereby NH_3_ is titrated to NH_4_^+^ (Wright and Wood, 2009; Wright et al., 1986; Zimmer, 2024). In alkaline water, when pH is near or exceeds the pK_a_ of the NH_3_ ←→ NH_4_^+^ equilibrium reaction (pK_a_ ∼ 9.0-9.5), this acid-trapping mechanism is inhibited which results in a reduction in ammonia excretion and an increase in the concentration of ammonia in the blood plasma and tissues (Chew et al., 2003; Kumai et al., 2015; Kwan et al., 2024; Laurent et al., 2000; McGeer and Eddy, 1998; Scott and Wilson, 2007; Scott et al., 2005; Thompson et al., 2016; Wilkie and Wood, 1991; Wilkie and Wood, 1994; Wilkie et al., 1993; Wilkie et al., 1996; Wilson et al., 1998; Wright et al., 1993; Yesaki and Iwama, 1992; Zhao et al., 2024; Zimmer and Perry, 2020; Zimmer et al., 2024). In addition, alkaline water bodies have been described as a “CO_2_ vacuum” (Johansen et al., 1975) which leads to a large outwardly directed P_CO2_ gradient such that fish exposed to alkaline conditions experience a respiratory alkalosis (McGeer and Eddy, 1998; Wilkie and Wood, 1991; Wilkie et al., 1996; Wright and Wood, 1985; Yesaki and Iwama, 1992). In some instances, pH can be maintained in the face of reduced plasma P_CO2_ via an increase in metabolic proton production (Wilkie et al., 1996), though this likely depends on water chemistry parameters such as [Ca^2+^] and [HCO_3_^-^] (Wilkie and Wood, 1996; Yesaki and Iwama, 1992). Fish exposed to high pH also experience a disruption in ion balance (Heming and Blumhagen, 1988; Kwan et al., 2024; McGeer and Eddy, 1998; Thompson et al., 2015; Wilkie and Wood, 1991; Wilkie et al., 1993; Yesaki and Iwama, 1992; Zimmer et al., 2024) due to the inhibitory effects of high pH on ion influx rates (Scott et al., 2005; Wilkie et al., 1999; Wright and Wood, 1985), although this effect is variable in some studies (Scott and Wilson, 2007). The mechanism underlying the inhibitory effect of high pH on ion uptake is currently not well understood but may involve a reduction in the availability of H^+^ and HCO_3_^-^ as counter ions for Na^+^ and Cl^-^ uptake, respectively, in response to respiratory alkalosis (Wilkie et al., 1999; Wood, 2022).

Despite the physiological challenges of life in alkaline waters, several examples exist of fish that inhabit waters of elevated pH. such as the Magadi tilapia, *Alcolapia grahami* (Randall et al., 1989), in Lake Magadi, Kenya (pH 9.6-10.0) and the tarek (pearl mullet), *Alburnus tarichi* (Danulat and Kempe, 1992), in Lake Van, Turkey (pH 9.8). More moderately alkaline lakes, however, can support diverse fish populations (McGeer et al., 1994; Mitchell and Prepas, 1990). The presence of fish species across these gradients of natural alkalinity and pH has been of great interest to the comparative physiology community given the considerable physiological challenges imposed by exposure to elevated alkalinity and pH.

Much of our understanding of the physiological responses to alkaline exposure has been generated from studying “alkaline-naïve” individuals. However, some fishes inhabiting naturally alkaline lakes have demonstrated increased alkaline tolerance (Wilkie and Wood, 1996; Wood, 2022). The most extreme example is the Magadi tilapia, which circumvents the inhibitory effect of its highly alkaline environment on ammonia excretion rates by excreting 100% of its nitrogenous waste as urea (Randall et al., 1989; Wood et al., 1989). Magadi tilapia express the full suite of ornithine urea cycle (OUC) enzymes as adults (Lindley et al., 1999; Randall et al., 1989), leading to a fully ureotelic physiology that appears to be obligatory (Wood, 2022). Other species inhabiting less alkaline environments, such as Pyramid Lake in Nevada, USA, show less dramatic adaptations including a reduction in nitrogen production and a shift to renal ammonia excretion (McGeer et al., 1994; Wilkie et al., 1993; Wilkie et al., 1994; Wright et al., 1993), and a tighter control of ion and acid-base balance in response to elevated pH (Wilkie et al., 1994). Tolerance of elevated plasma and tissue ammonia levels may also be critical to survival in alkaline waters in some species such as the naked/scaleless carp (*Gymnocpyris przewalskii*), which inhabits the alkaline Qinghai Lake in the Tibetan plateau (Wood, 2022). This species demonstrates much higher activities of glutamine synthetase and glutamate dehydrogenase, which catalyze ammonia-consuming reactions, compared to alkaline-naïve species (Wang et al., 2003; Wood, 2022). Moreover, several studies have identified a large number of genes related to nitrogen metabolism, ion regulation, acid-base regulation, and immune responses that appear to be under selection in species inhabiting alkaline lakes (Tong and Li, 2020; Tong et al., 2021; Xu et al., 2017; Zhang et al., 2015; Zhou et al., 2023).

The goal of the present study was to further characterize the mechanisms underlying alkaline tolerance in fish using a species collected from a naturally alkaline endorheic lake, Buffalo Lake (pH = 9.2), in Alberta, Canada. We collected brook stickleback (*Culaea inconstans*) from Buffalo Lake and a neutral reference lake (Buck Lake; pH = 8.2), and acclimated fish from both populations to common neutral lab conditions (pH = 8.0) for over 2 months. We hypothesized that Buffalo Lake stickleback have increased alkaline tolerance compared to Buck Lake stickleback and that this tolerance is a result of intraspecific differences in the physiological response to alkaline exposure. Based on this hypothesis, we predicted that in response to laboratory exposure to alkaline water (pH = 9.5) Buffalo Lake stickleback would exhibit smaller disruptions in nitrogen and ion balance compared to Buck Lake stickleback. Furthermore, based on previous evidence demonstrating the importance of branchial regulation of ion and nitrogen balance in response to alkaline exposure in other species (Laurent et al., 2000; Wilkie et al., 1994; Zhao et al., 2024), we predicted that changes in the activity of enzymes critical to nitrogen, ion, and acid-base regulation (Na^+^/K^+^-ATPase, H^+^-ATPase, and carbonic anhydrase) in response to alkaline exposure would differ between the populations and that this difference in activity would also be reflected by differences in transcriptomic responses in the gill. Overall, our results demonstrated clear intraspecific differences in alkaline tolerance between alkaline and neutral populations of brook stickleback, with these differences being at least partly explained by differences in ion and acid-base regulation and branchial gene expression.

## Materials and Methods

### Fish

Brook stickleback (*Culaea inconstans*) were collected from a neutral lake (Buck Lake; 52.992878, −114.714775; mean mass = 1.29 ± 0.05 g, range = 0.82 – 1.64 g) and an alkaline lake (Buffalo Lake; 52.530261, −112.815954; mean mass = 1.01 ± 0.04 g, range = 0.81 – 1.54 g) in Alberta, Canada [Alberta Fish Research Licence #21–3801] in September and October 2021 using standard minnow traps baited with commercial pet food. The chemical composition of Buffalo Lake and Buck Lake water are presented in Table 1. Fish were transferred from the traps to coolers containing aerated lake water held within 2 °C (temperature range at sites = 10 – 15 °C) of the water temperature at the site. Coolers were transported to the University of Alberta aquatics facilities and the fish were subsequently transferred to 100-L acrylic tanks filled with approximately 20 L of lake water. The tanks were then fitted with air stones and a supply of facility water (Table 1; 10-12 °C) at a rate of approximately 1 L min^-1^ such that the water in the tanks was gradually replaced with facility water. Stickleback from both populations were held in flow-through facility water (10-12 °C) under a 12:12 light:dark photoperiod and fed twice daily (10:00 and 15:00) to satiation with live brine shrimp. Mortality under these conditions was less than 5% overall for both populations. Fish were acclimated for at least 2 months prior to experimentation and were fasted for 24 h prior to all experimental procedures described below and remained fasted during experiments. Fish handling and experimentation was approved by the University of Alberta Biosciences Animal Care Committee (Animal Utilization Protocol 00003831).

**Table 1.** Chemical composition of lake water and lab exposure water. Values are in mmol L^-1^. Buffalo and Buck Lake data are presented as means of samples collected over 40 years of historical sampling with range included in brackets. Facility and alkaline water data are presented as means ± s.e.m of measurements made in the lab; for pH data, s.e.m. is replaced by the typical variation of the facility water and the titrator system that maintained pH in alkaline water.

|  | pH | [HCO<sub>3</sub><sup>-</sup>] | [Na<sup>+</sup>] | [Cl<sup>-</sup>] | [Ca<sup>2+</sup>] |
| --- | --- | --- | --- | --- | --- |
| **Buck Lake\*** | 8.20<br>(6.60 – 9.44)<br>n = 514 | 3.02<br>(0.07 – 3.04)<br>n = 45 | 0.65<br>(0.43 – 0.96)<br>n = 45 | 0.07<br>(0.03 – 0.14)<br>n = 44 | 0.65<br>(0.50 – 0.82)<br>n = 44 |
| **Buffalo Lake\*** | 9.19<br>(8.85 – 9.48)<br>n = 92 | 20.06<br>(13.79 – 20.58)<br>n = 92 | 23.65<br>(16.83 – 29.97)<br>n = 92 | 0.51<br>(0.31 – 0.96)<br>n = 92 | 0.21<br>(0.10 – 0.45)<br>n = 92 |
| **Facility water** | 8.0 ± 0.1 | 1.74 ± 0.13<br>n = 8 | 0.57 ± 0.08<br>n = 8 | 0.21 ± 0.01<br>n = 8 | 1.15 ± 0.12<br>n = 8 |
| **Alkaline water** | 9.5 ± 0.1 | 22.14 ± 0.30<br>n = 8 | 24.48 ± 0.66<br>n = 8 | 0.25 ± 0.02<br>n = 8 | 0.10 ± 0.01<br>n = 8 |

\*Data obtained from the Government of Alberta Water Quality Portal (https://environment.extranet.gov.ab.ca/apps/WaterQuality/dataportal/) accessed on July 16, 2024.

### Survival in alkaline water

Tolerance to alkaline water was tested in Buck Lake and Buffalo Lake stickleback. Alkaline water (Table 1) was achieved by adding 20 mmol L^-1^ NaHCO_3_ to facility water and adjusting pH to 9.5 using 1 M NaOH. Note that alkaline water was prepared at least 12 h prior to fish exposure to allow carbonate precipitates to settle out of solution. Fish were transferred in groups of 6-8 (n = 1) to 5-L static aquaria containing 4 L of alkaline water that were held in a 10-12 °C water bath. Aquaria were fitted with automatic titrators (pH Controller, BlueLab, New Zealand) that maintained pH at 9.5 ± 0.1 via the addition of 0.1 M NaOH. Water changes (80% of total volume) were conducted each day and fish were fasted for the duration of the 7-d exposure period. Throughout the exposure, fish were checked for signs of morbidity (primarily the loss of equilibrium), at which point they were euthanized using 1 g/L MS-222, and for cases of mortality, at which point they were removed from the tank. Daily morbidities and mortalities were recorded and used to determine survival rates in alkaline water. Note that for each population, one control survival experiment using normal facility water (pH = 8.0 ± 0.1) was conducted in order to ensure that mortality was not a function of the experimental procedure.

### Acute alkaline exposures

An additional set of experiments was conducted to determine population-specific effects of acute (1 or 4 d) exposure to alkaline water. For the 4-d experiment, fish were transferred individually to 200-mL black opaque plastic containers held in a 10-12 °C water bath. Each container was fitted with tubing that supplied the containers with recirculating facility water held at 10-12 °C and an air line to maintain constant aeration. Fish were held under these recirculating conditions overnight. The following morning, a 24-h control flux period in facility water was conducted. For this period, recirculating flow was stopped and water in the containers was lowered to a set volume (approximately 50 mL) and water samples (2 mL) were collected at 0 and 12 h, where thereafter water was replaced by gently flushing the containers with fresh facility water and samples were collected again at 0 and 12 h post-water change (i.e., 12-24 h of the control flux period). The containers were then flushed thoroughly with alkaline water (Table 1) that was prepared as described above. Complete turnover of the containers was monitored by a pH electrode that was placed in the container; the alkaline flux period began only once pH of the water was equal to 9.5. Water samples were collected every 12 h, with water changes occurring every 12 h following the procedure described above. Throughout the entire 4-d flux period, pH in the containers was maintained by automatic titration (pH Controller, BlueLab) with 0.01 M NaOH. At the end of the 4-d period, fish were removed from the containers and water volume was recorded. Fish were then euthanized using 1 g/L MS-222 titrated to pH 9.5 using NaOH, weighed, and the entire gill basket was excised, divided into two halves, flash frozen in liquid nitrogen, and stored at −80 °C. The viscera (gut, liver, kidney, and gonads) were removed, and the carcass of the fish was flash frozen in liquid nitrogen. To assess these lethal endpoints at 1 d of exposure, the same procedure described above was additionally conducted on fish exposed to alkaline water for 1 d, or to facility water for 1 d as a control group. Note that flux measurements were not conducted in these 1-d exposures.

### Analytical techniques and calculations

Water chemistry analysis was performed on laboratory waters (facility and alkaline water), whereas historical data were used to characterize the water composition of Buck Lake and Buffalo Lake (Table 1). Total CO_2_ in water samples was measured using a 965 CO_2_ analyzer (Corning, Corning, NY, USA) and [HCO_3_^-^] was calculated using the Henderson-Hasselbalch equation and αCO_2_ and pK_app_ values derived from Boutilier, Heming, & Iwama (1984). [Na^+^] and [Ca^2+^] were determined by atomic absorption flame spectroscopy (iCE 3500 AAS Atomic Absorption Spectrometer, ThermoFisher Scientific, Waltham, MA, USA) and [Cl^-^] was determined by a colourimetric assay (Zall et al., 1956).

[Ammonia] and [urea] in water samples were determined by colourimetric assays (Rahmatullah and Boyde, 1980; Verdouw et al., 1978). Ammonia and urea flux rates were calculated using the following equation:

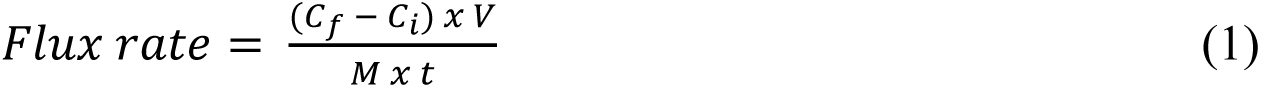

where C_f_ and C_i_ are the final and initial concentrations of ammonia-N or urea-N (µmol N L^-1^), V is volume (L), M is mass (g), and t is time (h).

Carcass analytes were determined following methods described previously (Zimmer et al., 2024). Briefly, frozen carcass samples were ground to a fine powder in a liquid nitrogen-cooled mortar and pestle. The powdered samples were deproteinized by the addition of 5 vols of 8% perchloric acid and incubation on ice for 5 min. Samples were then centrifuged for 2 min at 10 000 g to obtain a supernatant which was neutralized by the addition of 3 M KOH. [Ammonia] in the supernatant was determined using a commercial kit (Sigma-Aldrich, St. Louis, MO, USA), while [urea] was determined using the same colourimetric assay described above for water samples (Rahmatullah and Boyde, 1980). Similar to water samples, [Na^+^] and [Cl^-^] in neutralized tissue digests were measured by flame atomic absorption spectroscopy and a colourimetric assay (Zall et al., 1956), respectively, and [lactate] was determined by the lactate dehydrogenase method (Bergmeyer, 1983).

The activities of Na^+^/K^+^-ATPase, H^+^-ATPase, and carbonic anhydrase in gill tissue were assayed following established protocols (Henry, 1991; Lin and Randall, 1993; McCormick, 1993) described previously (Lim et al., 2015). Frozen gill tissue was homogenized on ice in a sodium deoxycholate buffer containing EGTA and centrifuged for 5 min at 10 000 g to remove cellular debris and cartilage. Na^+^/K^+^-ATPase and H^+^-ATPase activities in the collected supernatant (10 µL) were assayed indirectly by the rate of decrease in NADH concentration (measured at 340 nm using a plate spectrophotometer; SpectraMAX, Molecular Devices, Menlo Park, CA, USA) over a 15-min period in the presence or absence of ouabain (Na^+^/K^+^-ATPase inhibitor) or N-ethyl-maleimide (NEM) with sodium azide (H^+^-ATPase inhibitor). Activity of carbonic anhydrase in the gill homogenates (10 µL) was determined by the rate of pH change over a 30 s period following the addition of 1 mL CO_2_-saturated water to 10-mL reaction buffer (pH 7.4) held at 2-4 °C. Activity was determined by the difference between catalyzed (homogenate added) and non-catalyzed (no homogenate added) reaction rates. Protein concentration in the homogenates was determined using a commercial assay kit (Bio-Rad Protein Assay, Bio-Rad, Hercules, CA, USA). Na^+^/K^+^-ATPase and H^+^-ATPase activities were expressed as µmol ADP mg protein^-1^ h^-1^ and carbonic anhydrase activity was expressed as µmol H^+^ mg protein^-1^ min^-1^.

### Transcriptomic analyses

Frozen gill tissue (∼20-40 mg) was homogenized in 1 mL ice-cold Qiazol (Qiagen, Hilden, Germany) using a stator and rotor homogenizer. Total RNA was precipitated from the homogenate following the protocol of the manufacturer. RNA was reconstituted in nuclease-free water (20-50 µL) and RNA quantity and quality was assessed using a bio-analyzer (2100 Bioanalyzer, Agilent Technologies, Inc., Santa Clara, CA, USA). RNA integrity number (RIN) was > 9 in all samples. mRNA stranded library preparation (New England Biolabs, Ltd., Ipswich, MA, USA), quality control, and RNA sequencing (NovaSeq 6000 PE100, Illumina, Inc., San Diego, CA, USA) was performed by Genome Quebec (Montreal, QC, Canada). Quality of RNA reads was assessed using *FastQC* (v. 0.11.9; Brown *et al.,* 2017), and results were compiled with *MultiQC* (v.1.13; Ewels et al., 2016). Reads were then filtered and trimmed using *Trimmomatic* (v. 038; Bolger et al., 2014), with HEADCROP:12 applied after default treatment to account for remaining adapter sections after demultiplexing. Read quality was then reassessed using *FastQC* and *MultiQC*, which did not reveal any significant issues apart from high sequence duplication levels.

Trimmed reads were aligned to the three-spined stickleback (*Gasterosteus aculeatus*) reference genome (NCBI, University of Georgia, GCF_016920845.1, https://www.ncbi.nlm.nih.gov/datasets/genome/GCF_016920845.1/) using *STAR* (v. 2.7.7; Dobin et al., 2013). We selected this genome to align the reads because of its high quality annotation and because *C. inconstans* and this species only diverged about 25.7 million years ago (TimeTree; Kumar et al., 2022). The *outFilterScoreMinOverLRead* and *outFilterMatchNminOverLread* parameters were relaxed to 0.4 and 0.3 respectively, to account for nucleotide divergence between the genomes of *C. inconstans* and *G. aculeatus*. Read alignment was determined to be of excellent quality after sorting the reads with *samtools* (v. 1.11; Li et al., 2009). Sorted reads were quantified with *FeatureCounts* (v. 2.0.1) of the *Subread* package (Liao et al., 2014). Many of the reads could not be quantified because they were mapped to multiple loci on the genome, to regions on the genome not annotated in the NCBI annotations, or to multiple genes at once. The *FeatureCounts* counts matrix was pre-filtered using *DESeq2* following the developers’ recommendations (Love et al., 2014) by filtering out genes with less than 10 reads in the equivalent of one test group’s worth of samples (i.e., n = 6). Out of the 28,136 genes quantified using *FeatureCounts,* 8,200 genes were removed after pre-filtering due to low expression (Table S1). Notably, the number of quantified alignments was smaller than the number of raw reads for two reasons. First, many reads were removed from the data by STAR while mapping and aligning the reads (63.87% of total trimmed reads were mapped and aligned), most likely because of sequence divergence between the *C. inconstans* reads and the *G. aculeatus* reference genome. Second, *FeatureCounts* excludes multi-mapping reads (c. 418 million out of 1.44 billion alignments).

We applied five different Wald tests to the counts matrix using the DESeq() function. Two of them tested for differential gene expression caused by populational differences (“Buck” vs “Buffalo”) in fish exposed to one treatment or the other (“Treatment”), two others tested for differential gene expression caused by alkalinity exposure treatment (“Control” vs “Alkaline”) in both population groups (“Population”), and the fifth tested for the presence of an interaction between treatment and population. Comparisons for the Wald test were performed using the maximum likelihood estimators (MLEs) of normalized log2 average read counts for each of the groups. These MLEs were calculated using a negative binomial distribution constructed with the normalized log2 read counts. Due to the high number of tested genes, the Wald tests’ p-values were adjusted using the Benjamini-Hochberg method. For all tests, a gene was considered to have been differentially expressed due to a variable if the adjusted p-value of a gene’s Wald test for said variable was < 0.05, and if log2 fold change is > 2 (up-regulated) or < −2 (down-regulated).

A principal components analysis (PCA) was performed on the regularized-logarithm-transformed (*rlog*) (Love et al., 2014) read counts matrix with the plotPCA() function integrated in *BioConductor* (Huber et al., 2015). We also generated an UpSet plot (Lex et al., 2014) using the results of the genes’ Wald tests to observe in which ways differentially expressed genes were grouped according to their causes of differential expression. This was performed using the UpSet() function innate to the *ComplexHeatmap* package v. 2.22 (Gu et al., 2016), which uses code found in the *UpSetR* package (Conway et al., 2017). A gene clustering heatmap was also created using *ComplexHeatmap*’s pheatmap() function, which uses code from the *Pretty Heatmaps* package (Kolde, 2019). For this purpose, the *rlog*-transformed read counts matrix was filtered to include only the counts of genes that were differentially expressed. Counts for each gene were then centered and scaled to allow for better comparison of gene expression patterns between the genes, after which the values were used to select the colours of each square of the heatmap. Genes were then clustered using complete-linkage, hierarchical clustering according to differences between their expression patterns. To better understand why genes clustered the way they did in the heatmap, five additional binary heatmaps were generated and added to the right of the gene clustering heatmap which represent the results of each of the five Wald tests applied to the differentially expressed genes.

Version 3.19 of *goseq* was used to perform a gene ontology (GO) enrichment analysis on the reads (Young et al., 2010). Due to the nature of our dataset and the lack of GO annotations stored in the native *goseq* database, gene lengths and annotations had to be manually provided to the command. Gene lengths found in the *FeatureCounts* .csv output file were used to create a probability weighted function (PWF) using nullp() to determine the base probability of a gene being differentially expressed based on its length alone. The goseq() command was then used to perform GO enrichment analyses using information in the gene association file (GAF) linked to the *G. aculeatus* reference genome. The resulting tables were filtered to only include ontologies that were present in two or more of the differentially expressed genes.

### Statistics

Data for survival and physiological parameters are presented as means ± s.e.m. with individual data points presented as transparent symbols. Data points that were 1.5 times the interquartile range (IQR) below the 1^st^ quartile (Q1) or above the 3^rd^ quartile (Q3) were considered outliers and were omitted from the calculation of mean and s.e.m. and from statistical analyses; these outliers are represented as transparent symbols with a grey outline in plots. All statistical analyses and plotting were performed using R programming language (version 4.3.2) in RStudio (2023.12.1, Build 402; R Core Team 2023) and significance was accepted at the *P* < 0.05 level. Statistical significance was assessed using generalized linear models (GLMs) [glm() function; R Core Team 2023] with mass, lake origin, time in alkaline water (0 = control), and the interaction between lake origin and time as factors. Normality and heteroscedasticity were assessed visually using normal Q-Q plots and residual versus fitted values plots, respectively. Data were transformed (specific transformations are described in figure captions) if the model failed these qualitative visual inspections. The effects of mass, lake origin, time in alkaline water, and the interaction between lake origin and time were assessed using an ANOVA [aov() function; R Core Team 2023]. *P* values for each term and descriptions of data transformations are presented in corresponding figure captions. Estimated marginal means [emmeans() function; Lenth 2024] with a Tukey adjustment for multiple comparisons were used for post-hoc analyses. Effects of lake origin are represented by asterisks and effects of time in alkaline exposure are represented by uppercase letters for overall effects (i.e., lack of significant lake x time interaction term) and by lowercase letters for individual effects when the interaction term was significant.

## Results

Survival of Buck Lake and Buffalo Lake stickleback in holding conditions was high (> 95%) and survival over a 7-d period in control facility water (n = 1; data not shown) was 100%. Under alkaline conditions, Buffalo Lake stickleback showed 100% survival over a 7-d period, whereas survival in Buck Lake fish was significantly lower than that of Buffalo Lake fish by 4 d and declined significantly from the 0 d 100% survival value starting at 5 d of exposure (Fig. 1). At the termination of the 7-d exposure period, survival was only 14.6 ± 5.2 % in Buck Lake stickleback (Fig. 1).

**Figure 1.**
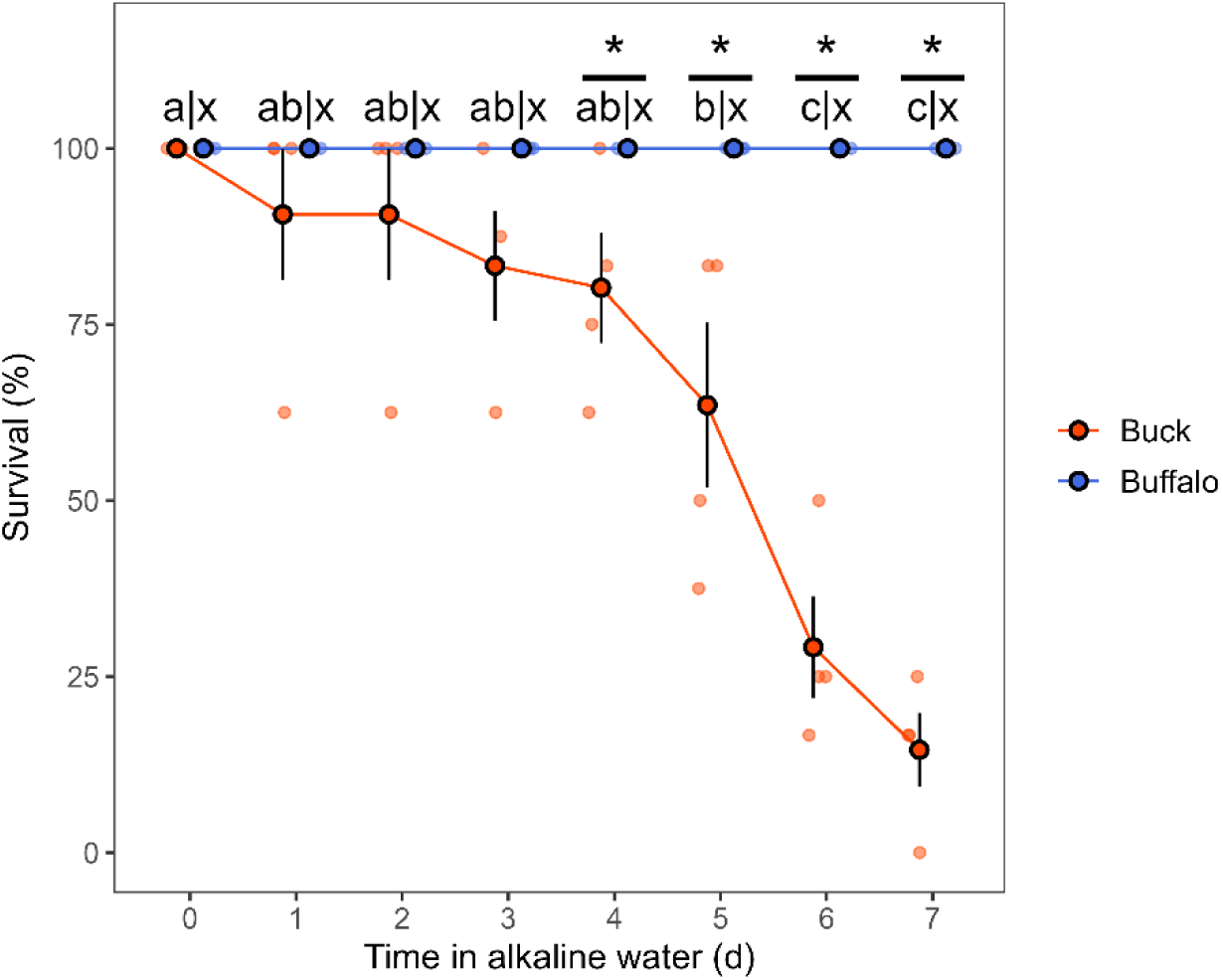
Survival of brook stickleback collected from Buck Lake (orange symbols) or Buffalo Lake (blue symbols) and exposed to alkaline water for 7 d. Asterisks represent time points at which there is a statistically significant effect of lake origin. Time points that do not share the same lowercase letter (abc = Buck Lake; xyz = Buffalo Lake) are significantly different from one another. [*P*_lake_ < 0.001, *P*_time_ < 0.001, *P*_lake*time_ < 0.001]

Ammonia excretion was not affected by lake origin and was initially inhibited by alkaline exposure but showed an overall recovery by 72-96 h of exposure (Fig. 2A). Urea excretion was significantly greater in Buffalo Lake stickleback compared to Buck Lake stickleback, but urea excretion was unaffected by the alkaline treatment (Fig. 2B). Tissue [ammonia] in carcass samples was not significantly different between Buffalo and Buck Lake fish, and significantly increased over the course of the 96-h exposure to alkaline water (Fig. 2C). Similar to urea excretion, tissue [urea] was significantly higher in Buffalo Lake stickleback, but no effects of time in alkaline water were observed (Fig. 2D).

**Figure 2.**
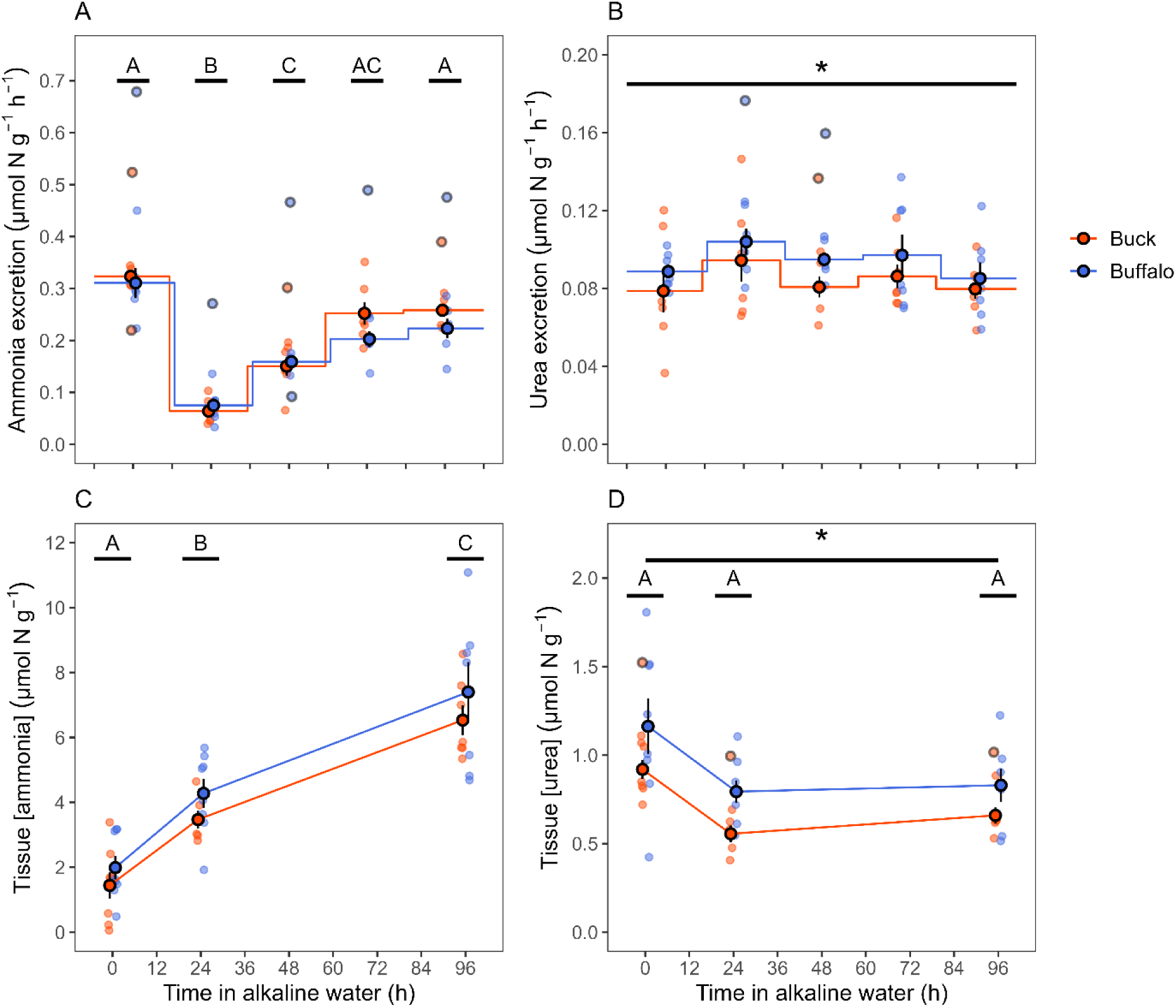
Ammonia-N (A) and urea-N (B) excretion rates and tissue [ammonia-N] (C) and [urea-N] (D) in carcass samples of brook stickleback collected from Buck Lake (orange symbols) or Buffalo Lake (blue symbols) and exposed to alkaline water for 96 h. A single asterisk represents an overall statistically significant effect of lake origin. Time points that do not share the same uppercase letter (ABC = overall effect of time) are significantly different from one another. [**A** – square root-transformed: *P*_mass_ = 0.585, *P*_lake_ = 0.848, *P*_time_ < 0.001, *P*_lake*time_ = 0.343; **B** – log-transformed: *P*_mass_ = 0.913, *P*_lake_ = 0.033, *P*_time_ = 0.256, *P*_lake*time_ = 0.954; **C** – square root-transformed: *P*_mass_ = 0.005, *P*_lake_ = 0.941, *P*_time_ < 0.001, *P*_lake*time_ = 0.931; **D** – log-transformed: *P*_mass_ = 0.935, *P*_lake_ = 0.015, *P*_time_ = 0.066, *P*_lake*time_ = 0.851]

There was no effect of lake origin on carcass tissue [Na^+^] which decreased significantly by 96 h of exposure to alkaline water relative to the levels observed in the 0 h control groups (Fig. 3A). Tissue [Cl^-^] also decreased over the duration of the alkaline exposure, however this effect was greater in Buck Lake stickleback in which tissue [Cl^-^] was significantly lower than that of Buffalo Lake fish at 24 and 96 h of exposure (Fig. 3B). Tissue lactate increased significantly by 96 h of exposure to alkaline water and showed no effects of lake origin (Fig. 3C). No effects of lake origin were observed for gill Na^+^/K^+^-ATPase, H^+^-ATPase, or carbonic anhydrase activity (Fig. 4). Gill H^+^-ATPase activity increased overall from 0 to 96 h of exposure to alkaline water (Fig. 4B).

**Figure 3.**
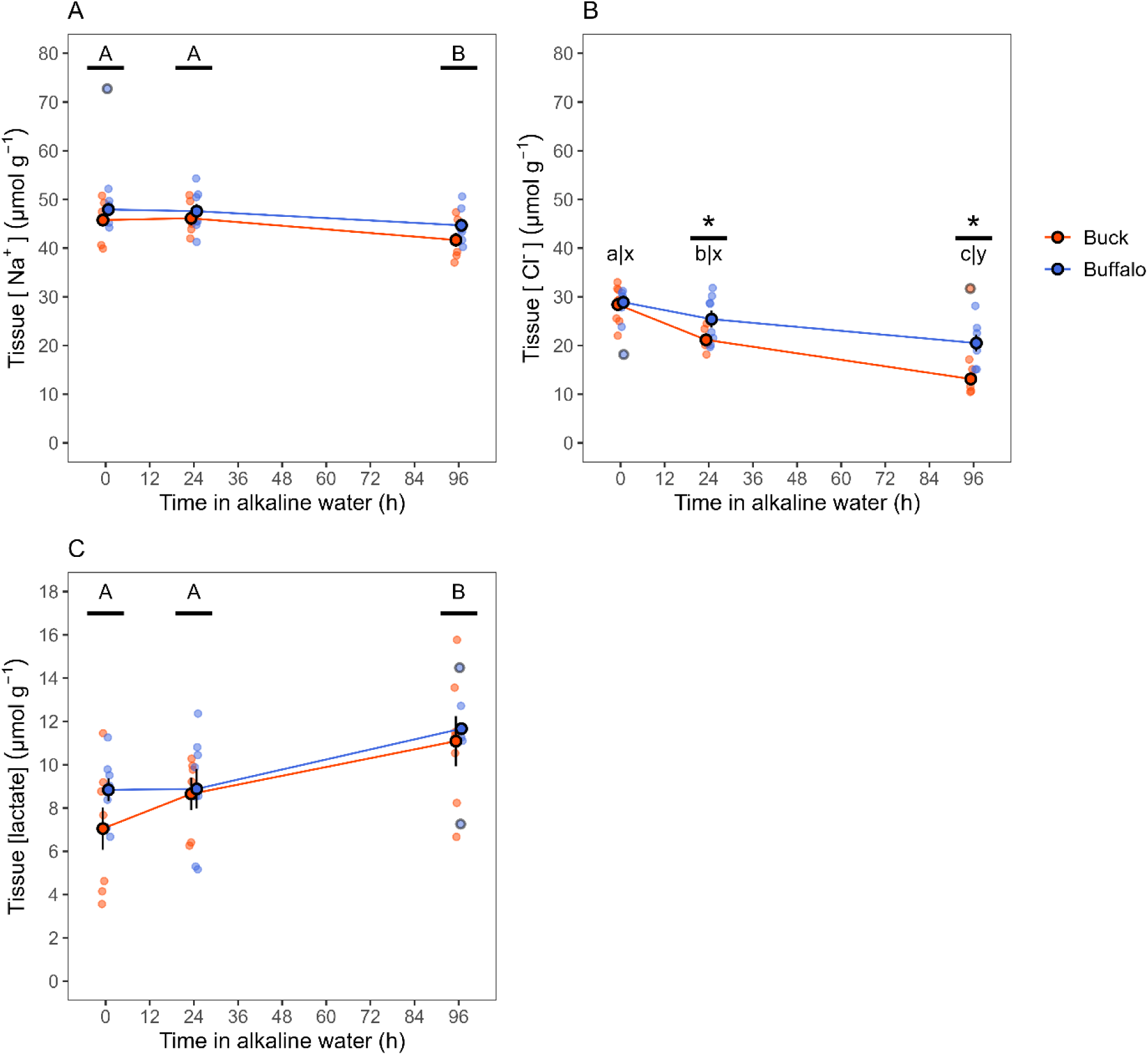
Tissue [Na^+^] (A), [Cl^-^] (B), and [lactate] (C) in carcass samples of brook stickleback collected from Buck Lake (orange symbols) or Buffalo Lake (blue symbols) and exposed to alkaline water for 96 h. Asterisks represent time points at which there is a statistically significant effect of lake origin. Time points within a population that do not share the same lowercase letter (abc = Buck Lake; xyz = Buffalo Lake) are significantly different from one another. Time points that do not share the same uppercase letter (ABC = overall effect of time) are significantly different from one another. [**A** – square-transformed: *P*_mass_ = 0.283, *P*_lake_ = 0.106, *P*_time_ = 0.018, *P*_lake*time_ = 0.914; **B** – log-transformed: *P*_mass_ = 0.651, *P*_lake_ < 0.001, *P*_time_ < 0.001, *P*_lake*time_ = 0.023; **C**: *P*_mass_ = 0.481, *P*_lake_ = 0.521, *P*_time_ = 0.001, *P*_lake*time_ = 0.654)]

**Figure 4.**
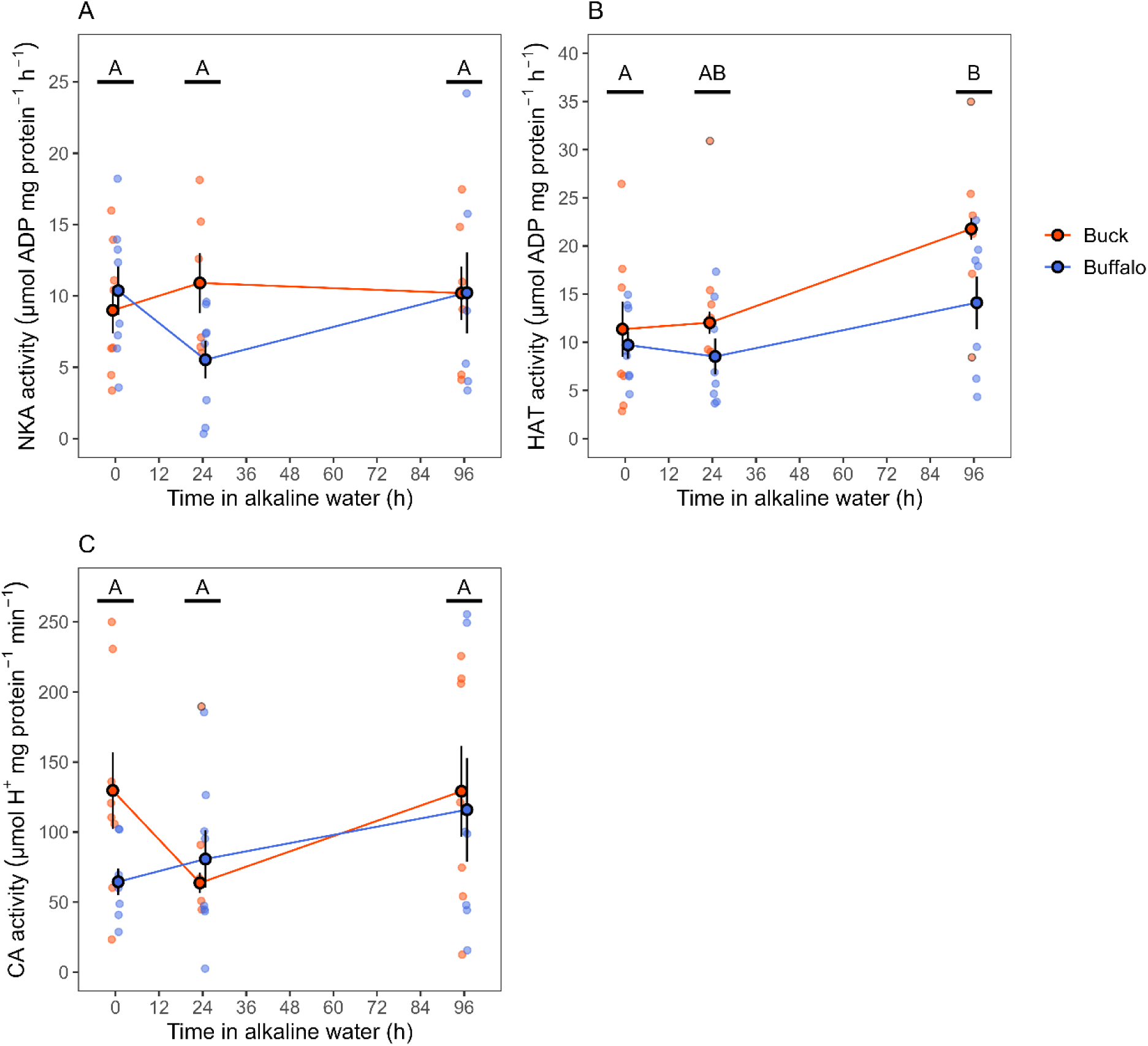
Activities of Na^+^/K^+^-ATPase (NKA; A), H^+^-ATPase (HAT; B), and carbonic anhydrase (CA; C) in gill tissue of brook stickleback collected from Buck Lake (orange symbols) or Buffalo Lake (blue symbols) and exposed to alkaline water for 96 h. Time points that do not share the same uppercase letter (ABC = overall effect of time) are significantly different from one another. [**A**: *P*_mass_ = 0.088, *P*_lake_ = 0.823, *P*_time_ = 0.744, *P*_lake*time_ = 0.174; **B**: *P*_mass_ = 0.001, *P*_lake_ = 0.756, *P*_time_ = 0.001, *P*_lake*time_ = 0.150; **C** – square-transformed: *P*_mass_ = 0.039, *P*_lake_ = 0.907, *P*_time_ = 0.221, *P*_lake*time_ = 0.341]

Differences in gene expression between populations and treatments were first examined by PCA (Fig. 5). PC1 accounted for 35% of the variation in gene expression among fish in our study and fish exposed to alkaline water appeared to increase along PC1 in principal component space. Notably, the change in PC1 was much greater for Buck Lake stickleback compared to Buffalo Lake stickleback. PC2 accounted for 14% of the variation and clearly delineated the two populations. These findings were corroborated by the results presented in the UpSet plot (Fig. 6) which depicts genes that were differentially expressed (488 total) in population and treatment comparisons and their interaction. The population and treatment combination that resulted in the largest number of differentially expressed genes was the effect of alkaline treatment in Buck Lake stickleback (267 genes total). Among these genes that were differentially expressed in Buck Lake stickleback in response to alkaline treatment, the majority were uniquely regulated by this treatment (147) or showed a population interaction (25). A small subset of these genes (35) was differentially expressed in both populations and additionally showed an interactive effect between populations (2). In contrast to Buck Lake fish, far fewer genes (100) showed differential regulation in response to alkaline treatment in Buffalo Lake stickleback. Similarly, fewer genes were differentially regulated between populations under control conditions (114 genes) compared to alkaline conditions (165 genes). Among these genes that were differentially expressed between populations, a total of 45 were differentially expressed irrespective of treatment.

**Figure 5.**
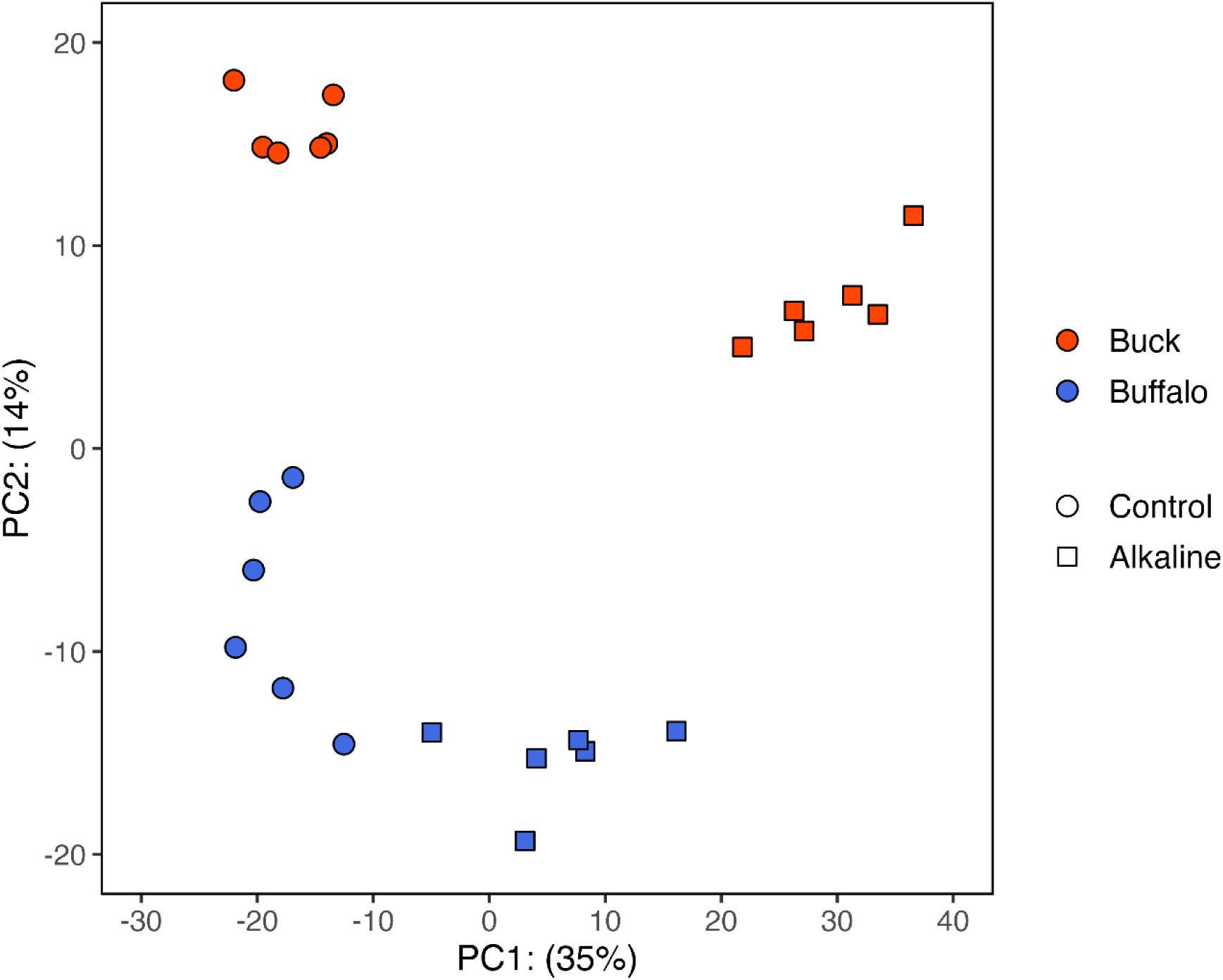
Principal component analysis (PCA) of genes expressed in gill tissue of brook stickleback collected from Buck Lake (orange symbols) or Buffalo Lake (blue symbols) and exposed to control (circles) or alkaline (squares) water for 96 h. PC1 accounted for 35% of the variation in gene expression and PC2 accounted for 14% of variation in gene expression.

**Figure 6.**
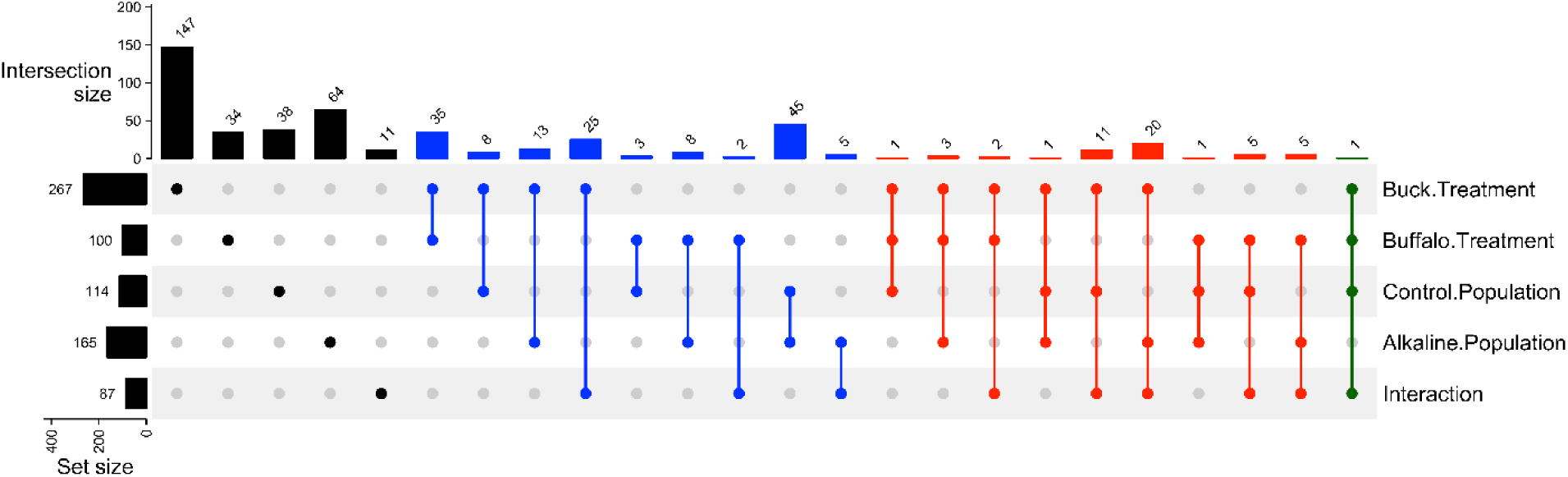
UpSet plot of differentially expressed genes in gill tissue of brook stickleback collected from Buck Lake or Buffalo Lake (“Population”) and exposed to control or alkaline water for 96 h (“Treatment”). The x-axis represents the number of differentially expressed genes within a given intersection of population and treatment combinations and their interaction. The y-axis represents the number of differentially expressed genes for a given population and treatment combination and their interaction.

The effect of population and treatment, and their interaction, on the expression of the 488 differentially expressed genes are also represented as a heat map plot (Fig. 7). This heat map analysis demonstrated 4 distinct clusters of genes with the clusters 1 and 3 representing genes that were downregulated or upregulated, respectively, in response to alkaline treatment. Similar to the PCA and UpSet plots, these heatmap clusters also clearly demonstrate that Buck Lake stickleback showed a much stronger gene expression response (both downregulation and upregulation in clusters 1 and 3, respectively) compared to Buffalo Lake stickleback. The other two clusters represent genes that were generally not affected by alkaline treatment but showed higher expression in Buffalo Lake fish (cluster 2) or showed higher expression in Buck Lake fish (cluster 4). The complete list of differentially expressed genes is included in a CSV file (Table S2) as supplementary material.

**Figure 7.**
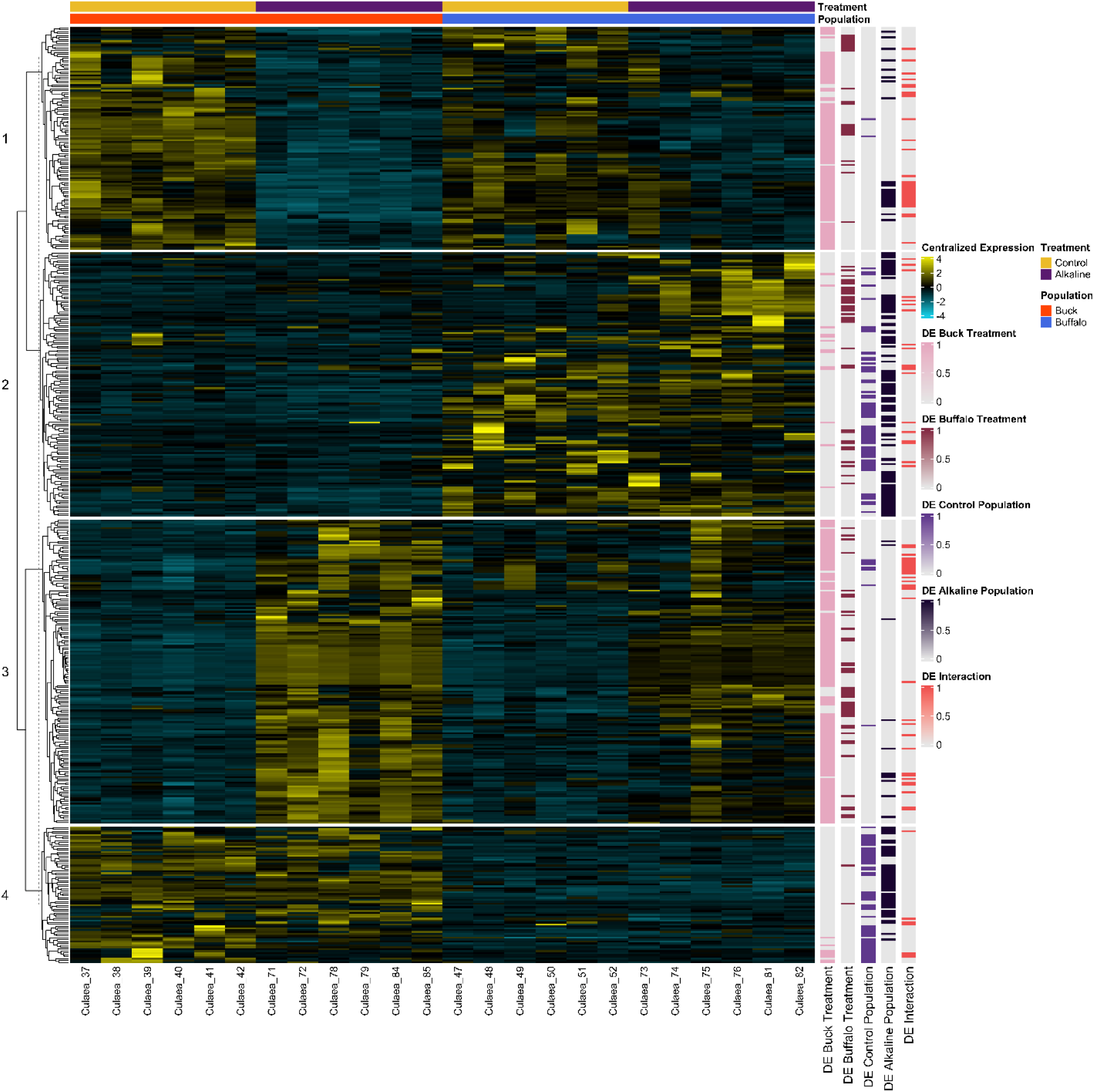
Gene clustering heatmap demonstrating centralized rlog-transformed read counts for genes that were differentially expressed between populations, in response to alkaline exposure, or due to an interaction effect. Individual fish are distributed across the x-axis of the map and are grouped according to population (Buck Lake = orange; Buffalo Lake = blue) and treatment (control = yellow; alkaline = purple). Each band on the heatmap represents a different gene; yellow bands represent higher expression, and blue bands represent lower expression. Binary heatmaps are included next to the expression heatmap that depict statistically significant differential gene expression (value of 1) for each population and treatment comparison.

GO enrichment analysis (Tables S3-S7) revealed relatively few terms related specifically to membrane transport functions that were over-expressed in our population and treatment comparisons, contrary to our initial hypothesis. However, we did observe some trends that may be related to intraspecific differences in alkaline tolerance between Buffalo Lake and Buck Lake stickleback. For instance, under control conditions, the term “membrane” consisted of 13 differentially regulated genes between populations, but this term was not significantly enriched (*P* = 0.069). Under alkaline conditions, however, “membrane” was the most over-represented GO term between populations (24 genes; *P* < 0.001). Moreover, in response to alkaline exposure, Buck Lake stickleback showed significant enrichment of several GO terms related to immune and stress responses including “immune response” (14 genes; *P* < 0.001), “positive regulation of T cell activation” (8 genes; *P* < 0.001), and “response to oxidative stress” (5 genes; *P* < 0.001).

In contrast, Buffalo Lake stickleback showed far fewer over-expressed GO terms in these categories in response to alkaline exposure. For example, the term “immune response” was non-significant (3 genes, *P* = 0.067).

## Discussion

The mechanisms used by fishes to tolerate and thrive in alkaline environments have been investigated in only a small number of highly specialized species, despite many alkaline lakes supporting diverse fish populations (Mitchell and Prepas, 1990; Wood, 2022). Therefore, we sought to characterize mechanisms of alkaline tolerance in brook stickleback inhabiting alkaline Buffalo Lake in Alberta, Canada. We initially hypothesized that brook stickleback from Buffalo Lake would have greater alkaline tolerance than those from a reference lake (Buck Lake), and that this would manifest as a tighter regulation of physiological processes that are typically impacted by exposure to alkaline or high pH water. Despite a 2-month acclimation to neutral holding conditions, Buffalo Lake stickleback showed no mortality when exposed to alkaline water of similar composition to Buffalo Lake (pH = 9.5) for a period of up to 7 d. In contrast, Buck Lake fish were unable to survive exposure to alkaline water that was similar in composition to Buffalo Lake water, demonstrating clear intraspecific differences in alkaline tolerance between these populations. By sampling fish at 4 d of exposure, prior to the observed decrease in survivorship in Buck Lake stickleback, we aimed to reveal the physiological basis for this lack of alkaline tolerance. Contrary to our hypothesis, we were able to detect only subtle differences in the physiological response to alkaline exposure between the two populations, with a reduction in Cl^-^ regulation and the implication of altered acid-base status (see *Ion and acid-base regulation* below), being the only intraspecific differences that we observed. In contrast, at the level of the transcriptome, there were very clear population-specific differences in gene expression patterns in the gill. Even under neutral control conditions, following 2 months of acclimation to common laboratory conditions, we observed a clear difference in the overall pattern of gene expression between populations that was largely associated with PC2 (14% of the total variation according to our PCA). Moreover, a much larger number of genes was differentially regulated in response to alkaline exposure in Buck Lake fish compared to those from Buffalo Lake. In Buck Lake stickleback, many of these genes were related to immune and stress responses, highlighting the fact that exposure to alkaline water is clearly much more stressful in this population. Overall, these findings highlight drastic intraspecific differences in alkaline tolerance that result from habitation of a naturally alkaline lake. Below we discuss these findings in the context of the physiological adaptations or adjustments that are necessary to cope with alkaline conditions.

### Nitrogen regulation

Based on our initial hypothesis that Buffalo Lake stickleback have increased alkaline tolerance due to the high pH of their environment (Table 1), we had predicted that they may show different patterns of nitrogen regulation in response to alkaline exposure compared to Buck Lake stickleback. Contrary to our hypothesis, both populations exhibited an initial inhibition of ammonia excretion rates upon exposure to alkaline water, followed by a recovery by 96 h of exposure (Fig. 2A), and there were no differences in carcass tissue [ammonia] between the populations (Fig. 2C). This pattern of ammonia excretion inhibition followed by a subsequent full or partial recovery has been demonstrated in many fish species exposed to alkaline conditions (Kumai et al., 2015; Scott et al., 2005; Wilkie and Wood, 1991; Wilkie et al., 1993; Wilkie et al., 1994; Wright et al., 1993). Recovery of ammonia excretion rates is likely a function of: (1) the accumulation of ammonia in the blood/tissues (Wilkie and Wood, 1991; Wilkie et al., 1993; Wilkie et al., 1994; Wright et al., 1993; Zhao et al., 2024) which, coupled to a blood alkalosis (see Introduction), presumably restores an outwardly directed P_NH3_ gradient; (2) changes in the function of branchial ionocytes (Laurent et al., 2000; Wilkie and Wood, 1994; Wilkie et al., 1994); and (3) increased expression of Rhesus (Rh) glycoproteins (Kumai et al., 2015; Sashaw et al., 2010; Zhao et al., 2024), which are ammonia channels that are critically important to the ammonia excretion mechanism of most fish species (Wright and Wood, 2009; Zimmer, 2024). In our transcriptomic analyses, the expression of one Rh gene, rhcga, was significantly increased in response to alkaline treatment in both populations, but there were no significant differences between populations (Table S2), indicating a potential for Rh-mediated ammonia excretion under alkaline conditions in this species. In zebrafish, rhcga (formerly Rhcg3) is not expressed in gill tissue (Nakada et al., 2007), which could indicate species-specific differences in Rh gene expression. In a comparison of genome-wide variation in single nucleotide polymorphisms (SNPs) between neutral and alkaline populations of Amur ide (*Leuciscus waleckii*), nonsynonymous SNPs in the coding region of rhcga were identified that appeared to be convergent across multiple species inhabiting alkaline environments (Zhou et al., 2023). Whether these same mutations exist in Buffalo Lake fishes, and the functional significance of these mutations in the context of alkaline physiology, are an interesting avenue for future research.

Among the physiological consequences of inhabiting high pH environments, the inhibition of ammonia excretion and ensuing accumulation of ammonia in the blood and tissues appears to be the most challenging (Wilkie and Wood, 1996; Wood, 2022). This challenge is perhaps best exemplified in the Magadi tilapia, which has adapted to its highly alkaline environment through a completely ureotelic nitrogen excretion strategy (Randall et al., 1989; Wood et al., 1989). In this species, urea is produced via the OUC, representing a pathway for scavenging ammonia and therefore preventing ammonia toxicity in an environment where ammonia excretion is virtually impossible due to the high pH and alkalinity that prevents NH_3_ diffusion at the gill (Wilkie and Wood, 1996; Wood, 2022). In species from less extreme alkaline environments, increased urea excretion and/or accumulation of urea in the plasma and tissues has also been observed (McGeer et al., 1994; Wilkie et al., 1993; Wright et al., 1993; Zhao et al., 2024), however this is generally believed to be a function of uricolysis because all of these species studied to date do not possess a functional OUC (McGeer et al., 1994; Wilkie et al., 1993; Wright et al., 1993). Similarly, brook stickleback collected from Buffalo Lake demonstrated an overall increase in both urea excretion rate (Fig. 2B) and carcass tissue [urea] (Fig. 2D), independent of alkaline exposure. The proportion of nitrogen excreted as urea, however, was not substantially different between Buck Lake and Buffalo Lake stickleback under alkaline conditions (23.4 ± 0.2 % and 25.7 ± 0.8 %, respectively, by 84-96 h of exposure). Therefore, it is unlikely that this increase production of urea contributes substantially to a reduction in the proportion of nitrogenous waste that must be excreted as ammonia. Notably, many alkaline resident species, including the Lahontan cutthroat trout (*Oncorhynchus clarkii henshawi*) from Pyramid Lake, show overall lower rates of ammonia production (McGeer et al., 1994; Wilkie et al., 1993; Wright et al., 1993) which have been implicated in their tolerance to alkaline conditions, however this did not appear to be the case for Buffalo Lake stickleback (Fig. 1). Therefore, we found little evidence to suggest that differences in the production or excretion of ammonia and urea are the underlying basis of increased alkaline tolerance in Buffalo Lake stickleback.

In contrast to the strategy of reduced ammonia production in Pyramid Lake species, other alkaline resident fish have adapted to alkaline conditions through increased tolerance of ammonia accumulation in the blood and tissues. The scaleless carp (*Gymnocypris przewalskii*) exhibits higher activities of ammonia-scavenging enzymes, glutamine synthetase and glutamate dehydrogenase, relative to alkaline-naïve species (Wang et al., 2003; Wood, 2022), and the gene coding for glutamine synthetase has been demonstrated to be under positive selection in this alkaline tolerant species (Zhang et al., 2015). Whether tissue-specific detoxification of ammonia is an important factor underlying increased alkaline tolerance of Buffalo Lake stickleback is currently unclear. Our transcriptomic analyses did not identify any differentially regulated genes that are putatively involved in nitrogen metabolism, however the gill is likely not an important site for ammonia detoxification compared to other organs. Indeed, at least one study has demonstrated that the number of differentially expressed genes in response to alkaline exposure in Amur ide resident to an alkaline lake was much greater in liver and kidney tissues compared to gill (Xu et al., 2013), suggesting that these tissues may play a more important in responding or adapting to alkaline conditions.

### Ion and acid-base regulation

Exposure to alkaline or high pH water often results in a disruption in ion and acid-base balance (Wilkie and Wood, 1996; Wood, 2022), and this was also observed in stickleback from both populations in the present study (Figs. 3A,B). Previous studies have attributed reductions in blood plasma and tissue [Na^+^] and [Cl^-^] to the inhibition of active ion uptake rates, with comparatively smaller effects on ion efflux rates (Scott et al., 2005; Wilkie et al., 1996; Wilkie et al., 1999). The mechanism of this inhibition has not been fully investigated, but has been attributed to a reduction in H^+^ and HCO_3_^-^ available for Na^+^/H^+^ and Cl^-^/HCO_3_^-^ exchanges in response to alkaline/high pH-induced respiratory alkalosis (Wilkie and Wood, 1991; Wilkie et al., 1999; Wood, 2022). In waters with high [HCO_3_^-^], inhibition of Cl^-^ uptake may additionally represent a direct inhibition of Cl^-^/HCO_3_^-^ exchange. Interestingly, the reduction in carcass tissue [Cl^-^] in response to alkaline exposure was greater in Buck Lake stickleback compared to Buffalo Lake stickleback (Fig. 3B), representing one of the few differences in the physiological response to alkaline water that was observed between the populations. This exacerbated response of tissue [Cl^-^] in Buck Lake fish may be indicative of differences in acid-base regulation since this response could result in an increase in the strong ion difference (SID; see Wood, 2022). In rainbow trout exposed to buffered water at pH 10.5, which was lethal by 3 d of exposure, trout experienced greater decreases in plasma [Cl^-^] compared to decreases in plasma [Na^+^] (i.e., increased SID) with a concomitant increase in [HCO_3_^-^] and pH (McGeer and Eddy, 1998), as would be predicted by SID theory. In contrast, in the alkaline-adapted Lahontan cutthroat trout, exposure to pH 10 in buffered Pyramid Lake water resulted in equal reductions of plasma [Na^+^] and [Cl^-^] such that SID was likely unchanged (Wilkie et al., 1993). In agreement, plasma [HCO_3_^-^] was unchanged and the alkalosis experienced by Lahontan cutthroat trout exposed to pH 10 was driven by a reduction in P_CO2_ (Wilkie et al., 1993). These results and those presented in our study suggest that regulation of SID in response to alkaline water may be an important aspect of adapting or acclimating to alkaline conditions. Although we were unable to directly assess acid-base status of stickleback exposed to alkaline water, the increase in carcass tissue [lactate], which has been observed in response to exposure to alkaline or high pH water in several species (Kwan et al., 2024; Wilkie and Wood, 1991; Wilkie et al., 1993; Wilson et al., 1998; Zimmer et al., 2024), would suggest an increase in the H^+^ production through increased glycolytic flux coupled to ATP hydrolysis (Hochachka and Mommsen, 1983), a process that would be necessary to counteract a respiratory alkalosis.

Ionoregulatory responses to alkaline water in rainbow trout (e.g., Wilkie et al., 1999) appear to be a function of changes in ionocyte density and surface area in the gill (Laurent et al., 2000; Wilkie and Wood, 1994), and similar changes occurred in Lahontan cutthroat trout exposed to Pyramid Lake water for up to 2 years (Wilkie et al., 1994). GO enrichment analysis revealed that under alkaline conditions the most over-expressed term between populations was “membrane” (Table S6), which was not significantly different under control conditions (Table S5). This might indicate that membrane reorganization or other membrane processes underlie some of the differences in alkaline tolerance between the populations. Moreover, branchial H^+^-ATPase increased in response to alkaline exposure, irrespective of population (Fig. 4), which might indicate differences in ionocyte function or density as a result of alkaline exposure, though would not explain differences in overall alkaline tolerance between populations. The role of H^+^-ATPase in ion and acid-base balance depends on its localization to the apical or basolateral membrane of ionocytes, which is a controversial topic in ionoregulatory physiology (Tseng et al., 2020; Tseng et al., 2022). In our study, the observed increase in gill H^+^-ATPase activity would make most sense in the context of basolateral expression which, in conjunction with increased metabolic H^+^ production, would contribute to the correction of a potential respiratory alkalosis in response to alkaline exposure. Interestingly, the expression of one H^+^-ATPase gene (V-type proton ATPase subunit d 1-like) showed a higher expression in Buffalo Lake fish compared to Buck Lake fish under control conditions, but was downregulated in response to alkaline exposure specifically in Buffalo Lake fish (Table S2). This discrepancy between gene expression and enzyme activity is not clear, but could be related to the fact that multiple genes encode different H^+^-ATPase subunits in fishes. Regardless, it is evident that exposure to alkaline water alters the expression of H^+^-ATPase in a potentially population-specific manner, but the role of this transporter in the physiological response to alkaline exposure requires further elucidation.

We identified a number of genes with putative ion and acid-base regulatory functions that were differentially expressed between populations or in response to alkaline treatment. Two carbonic anhydrase genes were differentially expressed in response to alkaline exposure specifically in Buffalo Lake fish. The expression of ca12 increased in response to alkaline exposure in this population and the expression of ca15b decreased in alkaline conditions (Table S2). At present, the cellular and subcellular localization of these carbonic anhydrase isoforms is not known in brook stickleback and therefore the implications of these results are unclear. In zebrafish, another ca15 isoform, ca15a, is believed to be expressed in the apical membrane of H^+^-ATPase-rich ionocytes (Gilmour and Perry, 2009; Lin et al., 2008), and its branchial expression is increased in response to acidic exposure (Lin et al., 2008). The reduction of an apical carbonic anhydrase isoform in response to alkaline exposure in Buffalo Lake stickleback would potentially act to retain H^+^ and additionally limit absorption of HCO_3_^-^ from the environment. In Amur ide, non-synonymous SNPs in ca15a and ca15b genes were found to be highly differentiated between alkaline and neutral populations (Zhou et al., 2023), suggesting that these genes may play important roles in the adaptation to alkaline environments. The gene encoding anion exchanger 2 (slc4a2a), which participates in Cl^-^/HCO_3_^-^ exchange, was upregulated in both populations in response to alkaline exposure, and this upregulation was more pronounced in Buffalo Lake stickleback (Table S2). Similar to our previous interpretations, it is difficult to surmise the significance of this response without an understanding of cellular and subcellular localization. It is clear, however, that relative to the Buck Lake stickleback from neutral lakes, Buffalo Lake fish appear to mount a greater transcriptional response to alkaline water with respect to genes with putative ion and acid-base regulatory functions, which could contribute to the increased tolerance of Buffalo Lake fish to alkaline conditions and the observed intraspecific differences in the regulation Cl^-^ balance.

### Conclusions and perspectives

The present study on brook stickleback from a naturally alkaline lake adds to the body of literature demonstrating increased alkaline tolerance in many alkaline resident fish species (Danulat, 1995; Wilkie and Wood, 1996; Wood, 2022). In some instances, this increased tolerance appears to be evolved through selection (Tong et al., 2021; Zhang et al., 2015; Zhou et al., 2023), however we cannot discern whether this is the case for brook stickleback in our study. The retention of increased alkaline tolerance after 2 months of acclimation to neutral laboratory conditions in Buffalo Lake stickleback could potentially be a function of developmental plasticity and common-garden experiments would be necessary to determine whether alkaline tolerance in this population is a function of local adaptation. What is clear is that this population represents physiological diversity, and it would be interesting to determine whether other species in Buffalo Lake and the many other alkaline lakes in Alberta, Canada demonstrate similar increases in alkaline tolerance relative to neutral lake populations. If this is a widespread phenomenon, it may be necessary to reevaluate conservation management approaches in these lakes because stocking efforts in these lakes may prove difficult given that resident populations may be uniquely adapted or acclimated to alkaline conditions. For example, high alkalinity has been identified as a potential challenge to stocking of some lakes in British Columbia, Canada (Thompson et al., 2015). Therefore, a better understanding of the potentially subtle mechanisms underlying increased alkaline tolerance in fish resident to alkaline lakes is an important task moving forward that will assist in designing effective management approaches. Moreover, this research adds to our fundamental understanding of the physiological processes underlying responses to alkaline exposure which has important implications for predicting the effects of transient alkalinization events such ash deposition from wildfires (Kwan et al., 2024) and eutrophication (Scott et al., 2005).

## Notes

### Competing Interest Statement

The authors have declared no competing interest.

## References

Bergmeyer, H. (1983). Methods of enzymatic analysis. New York: Academic Press.

Bolger, A. M., Lohse, M. and Usadel, B. (2014). Trimmomatic: A flexible trimmer for Illumina sequence data. Bioinformatics 30, 2114–2120.

Boutilier, R. G., Heming, T. A. and Iwama, G. K. (1984). Physiochemical parameters for use in fish respiratory physiology. In Fish Physiology (Vol. 10) (ed. Hoar, W.) and Randall, D. J.), pp. 403–430. New York: Academic Press.

Brauner, C. J., Gonzalez, R. J. and Wilson, J. M. (2013). Extreme environments: hypersaline, alkaline, and ion-poor waters. In Fish Physiology (Vol. 32) (ed. Mccorrnick, S. D.), Farrell, A. P.), and Brauner, C. J.), pp. 435–476. New York: Academic Press.

Chew, S. F., Hong, L. N., Wilson, J. M., Randall, D. J. and Ip, Y. K. (2003). Alkaline environmental pH has no effect on ammonia excretion in the mudskipper *Periophthalmodon schlosseri* but inhibits ammonia excretion in the related species *Boleophthalmus boddaerti*. Physiol. Biochem. Zool. 76, 204–214.

Conway, J. R., Lex, A. and Gehlenborg, N. (2017). UpSetR: An R package for the visualization of intersecting sets and their properties. Bioinformatics 33, 2938–2940.

Danulat, E. (1995). Biochemical-physiological adaptations of teleosts to highly alkaline, saline lakes. In Biochemistry and Molecular Biology of Fishes (ed. Mommsen, T. P.) and Hochachka, P. W.), pp. 229–249. Amsterdam: Elsevier.

Dobin, A., Davis, C. A., Schlesinger, F., Drenkow, J., Zaleski, C., Jha, S., Batut, P., Chaisson, M. and Gingeras, T. R. (2013). STAR: Ultrafast universal RNA-seq aligner. Bioinformatics 29, 15–21.

Ewels, P., Magnusson, M., Lundin, S. and Käller, M. (2016). MultiQC: Summarize analysis results for multiple tools and samples in a single report. Bioinformatics 32, 3047–3048.

Gilmour, K. M. and Perry, S. F. (2009). Carbonic anhydrase and acid-base regulation in fish. J. Exp. Biol. 212, 1647–1661.

Gu, Z., Eils, R. and Schlesner, M. (2016). Complex heatmaps reveal patterns and correlations in multidimensional genomic data. Bioinformatics 32, 2847–2849.

Heming, T. A. and Blumhagen, K. A. (1988). Plasma acid-base and electrolyte states of rainbow trout exposed to alum (aluminum sulphate) in acidic and alkaline environments. Aquat. Toxicol. 12, 125–139.

Henry, R. P. (1991). Techniques for measuring carbonic anhydrase activity in vitro: the electrometric delta pH and pH stat assay. In The Carbonic Anhydrases: Cellular Physiology and Genetics (ed. Dodgson, S. J.), E, T. R.), Gros, G.), and D, C. N.), pp. 119–126. New York: Plenum.

Hochachka, P. W. and Mommsen, T. P. (1983). Protons and anaerobiosis. Science 219, 1391–1397.

Huber, W., Carey, V. J., Gentleman, R., Anders, S., Carlson, M., Carvalho, B. S., Bravo, H. C., Davis, S., Gatto, L., Girke, T., et al. (2015). Orchestrating high-throughput genomic analysis with Bioconductor. Nat. Methods 12, 115–121.

Johansen, K., Maloiy, G. M. O. and Lykkeboe, G. (1975). A fish in extreme alkalinity. Respir. Physiol. 24, 159–162.

Kolde, R. (2019). Package: Pretty Heatmaps (pheatmap) 1.0.12.

Kumai, Y., Harris, J., Al-Rewashdy, H., Kwong, R. W. M. and Perry, S. F. (2015). Nitrogenous waste handling by larval zebrafish *Danio rerio* in alkaline water. Physiol. Biochem. Zool. 88, 137–145.

Kumar, S., Suleski, M., Craig, J. M., Kasprowicz, A. E., Sanderford, M., Li, M., Stecher, G. and Hedges, S. B. (2022). TimeTree 5: An expanded resource for species divergence times. Mol. Biol. Evol. 39, msac174.

Kwan, G. T., Sanders, T., Huang, S., Kilaghbian, K., Sam, C., Wang, J., Weihrauch, K., Wilson, R. W. and Fangue, N. A. (2024). Impacts of ash-induced environmental alkalinization on fish physiology, and their implications to wildfire-scarred watersheds. Sci. Total Environ. 953, 176040.

Laurent, P., Wilkie, M. P., Chevalier, C. and Wood, C. M. (2000). The effect of highly alkaline water (pH 9.5) on the morphology and morphometry of chloride cells and pavement cells in the gills of the freshwater rainbow trout: Relationship to ionic transport and ammonia excretion. Can. J. Zool. 78, 307–319.

Lenth, R. V (2024). emmeans: Estimated marginal means, aka least-squares means.

Lex, A., Gehlenborg, N., Strobelt, H., Vuillemot, R. and Pfister, H. (2014). UpSet: Visualization of intersecting sets. IEEE Trans. Vis. Comput. Graph. 20, 1983–1992.

Li, H., Handsaker, B., Wysoker, A., Fennell, T., Ruan, J., Homer, N., Marth, G., Abecasis, G. and Durbin, R. (2009). The Sequence Alignment/Map format and SAMtools. Bioinformatics 25, 2078–2079.

Liao, Y., Smyth, G. K. and Shi, W. (2014). FeatureCounts: An efficient general purpose program for assigning sequence reads to genomic features. Bioinformatics 30, 923–930.

Lim, M. Y. T., Zimmer, A. M. and Wood, C. M. (2015). Acute exposure to waterborne copper inhibits both the excretion and uptake of ammonia in freshwater rainbow trout (*Oncorhynchus mykiss*). Comp. Biochem. Physiol. C 168, 48–54.

Lin, H. and Randall, D. J. (1993). H^+^-ATPase activity in crude homogenates of fish gill tissue: inhibitor sensitivity and environmental and hormonal regulation. J. Exp. Biol. 180, 163–174.

Lin, T. Y., Liao, B. K., Horng, J. L., Yan, J. J., Hsiao, C. Der and Hwang, P. P. (2008). Carbonic anhydrase 2-like a and 15a are involved in acid-base regulation and Na^+^ uptake in zebrafish H^+^-ATPase-rich cells. Am. J. Physiol. Cell Physiol. 294, C1250–C1260.

Lindley, T. E., Scheiderer, C. L., Walsh, P. J., Wood, C. M., Bergman, H. L., Bergman, A. L., Laurent, P., Wilson, P. and Anderson, P. M. (1999). Muscle as the primary site of urea cycle enzyme activity in an alkaline lake-adapted tilapia, *Oreochromis alcalicus grahami*. J. Biol. Chem. 274, 29858–29861.

Love, M. I., Huber, W. and Anders, S. (2014). Moderated estimation of fold change and dispersion for RNA-seq data with DESeq2. Genome Biol. 15, 1–21.

McCormick, S. D. (1993). Methods for nonlethal gill biopsy and measurement of Na^+^, K^+^-ATPase activity. Can. J. Aquat. Sci. 50, 656–658.

McGeer, J. C. and Eddy, F. B. (1998). Ionic regulation and nitrogenous excretion in rainbow trout exposed to buffered and unbuffered freshwater of pH 10.5. Physiol. Zool. 71, 179–190.

McGeer, J. C., Wright, P. A., Wood, C. M., Wilkie, M. P., Mazur, C. F. and Iwama, G. K. (1994). Nitrogen excretion in four species of fish from an alkaline lake. Trans. Am. Fish. Soc. 123, 824–829.

Mitchell, P. and Prepas, E. E. (1990). Atlas of Alberta Lakes. University of Alberta Press.

Nakada, T., Hoshijima, K., Esaki, M., Nagayoshi, S., Kawakami, K. and Hirose, S. (2007). Localization of ammonia transporter Rhcg1 in mitochondrion-rich cells of yolk sac, gill, and kidney of zebrafish and its ionic strength-dependent expression. Am. J. Physiol. Regul. Integr. Comp. Physiol. 293, 1743–1753.

Pinheiro, J. P. S., Windsor, F. M., Wilson, R. W. and Tyler, C. R. (2021). Global variation in freshwater physico-chemistry and its influence on chemical toxicity in aquatic wildlife. Biol. Rev. 96, 1528–1546.

R Core Team (2024). R: a language and environment for statistical computing. Vienna: R Foundation for Statistical Computing.

Rahmatullah, M. and Boyde, T. R. C. (1980). Improvements in the determination of urea using diacetyl monoxime; methods with and without deproteinisation. Clin. Chim. Acta 107, 3–9.

Randall, D. J., Wood, C. M., Perry, S. F., Bergman, H., Maloiy, G. M. O., Mommsen, T. P. and Wright, P. A. (1989). Urea excretion as a strategy for survival in a fish living in a very alkaline environment. Nature 337, 165–166.

Sashaw, J., Nawata, M., Thompson, S., Wood, C. M. and Wright, P. A. (2010). Rhesus glycoprotein and urea transporter genes in rainbow trout embryos are upregulated in response to alkaline water (pH 9.7) but not elevated water ammonia. Aquat. Toxicol. 96, 308–313.

Scott, D. M. and Wilson, R. W. (2007). Three species of fishes from an eutrophic, seasonally alkaline lake are not more tolerant to acute exposure to high pH in the laboratory. J. Fish Biol. 70, 551–566.

Scott, D. M., Lucas, M. C. and Wilson, R. W. (2005). The effect of high pH on ion balance, nitrogen excretion and behaviour in freshwater fish from an eutrophic lake: A laboratory and field study. Aquat. Toxicol. 73, 31–43.

Thompson, W. A., Rodela, T. M. and Richards, J. G. (2015). The effects of strain and ploidy on the physiological responses of rainbow trout (*Oncorhynchus mykiss*) to pH 9.5 exposure. Comp. Biochem. Physiol. B 183, 22–29.

Thompson, W. A., Rodela, T. M. and Richards, J. G. (2016). Hardness does not affect the physiological responses of wild and domestic strains of diploid and triploid rainbow trout *Oncorhynchus mykiss* to short-term exposure to pH 9.5. J. Fish Biol. 89, 1345–1358.

Tong, C. and Li, M. (2020). Genomic signature of accelerated evolution in a saline-alkaline lake-dwelling Schizothoracine fish. Int. J. Biol. Macromol. 149, 341–347.

Tong, C., Li, M., Tang, Y. and Zhao, K. (2021). Genomic signatures of shifts in selection and alkaline adaptation in highland fish. Genome Biol. Evol. 13, evab086.

Tseng, Y. C., Yan, J. J., Furukawa, F. and Hwang, P. P. (2020). Did acidic stress resistance in vertebrates evolve as Na^+^/H^+^ exchanger-mediated ammonia excretion in fish? BioEssays 42, 1–7.

Tseng, Y. C., Yan, J. J., Furukawa, F., Chen, R. D., Lee, J. R., Tsou, Y. L., Liu, T. Y., Tang, Y. H. and Hwang, P. P. (2022). Teleostean fishes may have developed an efficient Na^+^ uptake for adaptation to the freshwater system. Front. Physiol. 13, 1–15.

Verdouw, H., Van Echteld, C. J. A. and Dekkers, E. M. J. (1978). Ammonia determination based on indophenol formation with sodium salicylate. Water Res. 12, 399–402.

Wang, Y. S., Gonzalez, R. J., Patrick, M. L., Grosell, M., Zhang, C., Feng, Q., Du, J. Z., Walsh, P. J. and Wood, C. M. (2003). Unusual physiology of scale-less carp, Gymnocypris przewalskii, in Lake Qinghai: A high altitude alkaline saline lake. Comp. Biochem. Physiol. A 134, 409–421.

Wilkie, M. P. and Wood, C. M. (1991). Nitrogenous waste excretion, acid-base regulation, and ionoregulation in rainbow trout (*Oncorhynchus mykiss*) exposed to extremely alkaline water. Physiol. Zool. 64, 1069–1086.

Wilkie, M. P. and Wood, C. M. (1994). The effects of extremely alkaline water (pH 9.5) on rainbow trout gill function and morphology. J. Fish Biol. 45, 87–98.

Wilkie, M. P. and Wood, C. M. (1996). The adaptations of fish to extremely alkaline environments. Comp. Biochem. Physiol. B 113, 665–673.

Wilkie, M. P., Wright, P. A., Iwama, G. K. and Wood, C. M. (1993). The physiological responses of the Lahontan cutthroat trout (*Oncorhynchus clarki henshawi*), a resident of highly alkaline Pyramid Lake (pH 9.4), to challenge at pH 10. J. Exp. Biol. 175, 173–194.

Wilkie, M. P., Wright, P. A., Iwama, G. K. and Wood, C. M. (1994). The physiological adaptations of the Lahontan cutthroat trout (*Oncorhynchus clarki henshawi*) following transfer from well water to the highly alkaline waters of Pyramid Lake, Nevada (pH 9.4). Physiol. Zool. 67, 355–380.

Wilkie, M. P., Simmons, H. E. and Wood, C. M. (1996). Physiological adaptations of rainbow trout to chronically elevated water pH (pH = 9.5). J. Exp. Zool. 274, 1–14.

Wilkie, M. P., Laurent, P. and Wood, C. M. (1999). The physiological basis for altered Na^+^ and Cl^-^ movements across the gills of rainbow trout (*Oncorhynchus mykiss*) in alkaline (pH = 9.5) water. Physiol. Biochem. Zool. 72, 360–368.

Wilson, J. M., Iwata, K., Iwama, G. K. and Randall, D. J. (1998). Inhibition of ammonia excretion and production in rainbow trout during severe alkaline exposure. Comp. Biochem. Physiol. B 121, 99–109.

Wood, C. M. (1993). Ammonia and urea metabolism and excretion. In The physiology of fishes (ed. Evans, D. H.), pp. 379–425. Boca Raton: CRC Press.

Wood, C. M. (2022). Conservation aspects of osmotic, acid-base, and nitrogen homeostasis in fish. In Fish Physiology (Vol. 39A) (ed. Cooke, S. J.), Fangue, N. A.), Farrell, A. P.), Brauner, C. J.), and Eliason, E. J.), pp. 321–388. Academic Press.

Wood, C. M., Perry, S. F., Wright, P. A., Bergman, H. L. and Randall, D. J. (1989). Ammonia and urea dynamics in the Lake Magadi tilapia, a ureotelic teleost fish adapted to an extremely alkaline environment. Respir. Physiol. 77, 1–20.

Wright, P. A. and Wood, C. M. (1985). An analysis of branchial ammonia excretion in the freshwater rainbow trout: Effects of environmental pH change and sodium uptake blockade. J. Exp. Biol. 114, 329–353.

Wright, P. A. and Wood, C. M. (2009). A new paradigm for ammonia excretion in aquatic animals: Role of rhesus (Rh) glycoproteins. J. Exp. Biol. 212, 2303–2312.

Wright, P., Heming, T. and Randall, D. (1986). Downstream pH changes in water flowing over the gills of rainbow trout. J. Exp. Biol. 126, 499–512.

Wright, P. A., Iwama, G. K. and Wood, C. M. (1993). Ammonia and urea excretion in Lahontan cutthroat trout (*Oncorhynchus clarki henshawi*) adapted to the highly alkaline Pyramid Lake (pH 9.4). J. Exp. Biol. 175, 153–172.

Xu, J., Li, Q., Xu, L., Wang, S., Jiang, Y., Zhao, Z., Zhang, Y., Li, J., Dong, C., Xu, P., et al. (2013). Gene expression changes leading extreme alkaline tolerance in Amur ide (*Leuciscus waleckii*) inhabiting soda lake. BMC Genomics 14, 682.

Xu, J., Li, J. T., Jiang, Y., Peng, W., Yao, Z., Chen, B., Jiang, L., Feng, J., Ji, P., Liu, G., et al. (2017). Genomic basis of adaptive evolution: The survival of amur ide (*Leuciscus waleckii*) in an extremely alkaline environment. Mol. Biol. Evol. 34, 145–159.

Yesaki, T. Y. and Iwama, G. K. (1992). Survival, acid-base regulation, ion regulation, and ammonia excretion in rainbow trout in highly alkaline hard water. Physiol. Zool. 65, 763–787.

Young, M. D., Wakefield, M. J., Smyth, G. K. and Oshlack, A. (2010). Gene ontology analysis for RNA-seq: accounting for selection bias. Genome Biol. 11, R14.

Zall, D. M., Fisher, D. and Garner, M. Q. (1956). Photometric determination of chlorides in water. Anal. Chem. 28, 1665–1668.

Zhang, R., Ludwig, A., Zhang, C., Tong, C., Li, G., Tang, Y., Peng, Z. and Zhao, K. (2015). Local adaptation of *Gymnocypris przewalskii* (Cyprinidae) on the Tibetan Plateau. Sci. Rep. 5, 09780.

Zhao, X. F., Huang, J., Li, W., Wang, S. Y., Liang, L. Q., Zhang, L. M., Liew, H. J. and Chang, Y. M. (2024). Rh proteins and H^+^ transporters involved in ammonia excretion in Amur Ide (*Leuciscus waleckii*) under high alkali exposure. Ecotoxicol. Environ. Saf. 273, 116160.

Zhou, Z., Yang, J., Lv, H., Zhou, T., Zhao, J., Bai, H., Pu, F. and Xu, P. (2023). The adaptive evolution of *Leuciscus waleckii* in Lake Dali Nur and convergent evolution of cypriniformes fishes inhabiting extremely alkaline environments. Genome Biol. Evol. 15, evad082.

Zimmer, A. M. (2024). Ammonia excretion by the fish gill: discoveries and ideas that shaped our current understanding. J. Comp. Physiol. B 194, 697–715.

Zimmer, A. M. and Perry, S. F. (2020). The Rhesus glycoprotein Rhcgb is expendable for ammonia excretion and Na^+^ uptake in zebrafish (*Danio rerio*). Comp. Biochem. Physiol. A 247, 110722.

Zimmer, A. M., Woods, O., Glover, C. N. and Goss, G. G. (2024). Exposure to alkaline water reduces thermal tolerance, but not thermal plasticity, in brook stickleback (*Culaea inconstans*) collected from an alkaline lake. Hydrobiologia 851, 2641–2655.

